# Relationship between ecological stoichiometry and community diversity of plant ecosystems in the upper reaches of the Tarim River, China

**DOI:** 10.1101/432278

**Authors:** Jingjing Zhao, Lu Gong, Xin Chen

## Abstract

**Aim:** Although it is commonly proposed that nutrient cycling can impact plant community diversity, this relationship has not been fully examined in arid and semi-arid zones. Here, we expand on the framework for evaluating the relationship between biodiversity and ecological stoichiometry by scaling up from the level of the community.

**Location:** The upper reaches of the Tarim River (Northwest China, 80°10’-84°36’E, 0°25’-41°10’N).

**Methods:** We used multivariate analysis of variance to compare the stoichiometric characteristics and species diversity indices of sampled plant communities. We also measured carbon (C), nitrogen (N), and phosphorous (P) content of plants. We then assessed correlations between community stoichiometry and species diversity through structural equation models (SEM) and redundancy analysis (RDA).

**Results:** We found that the differences between stoichiometric characteristics and community diversity indices were highly significant. The Margalef index was strongly related to C and P content. The Simpson’s index and Shannon-Weaner index were most strongly correlated with C content. Pielou’s index was closely related to C and N contents, and the C:N and C:P ratios were important at driving ecological dominance.

**Main Conclusions:** Our study highlights the importance of ecological stoichiometry in driving community assembly and diversity within a desert ecosystems in northwestern China. The relationship between eclogical stoichiometry in the desert plant community had an effect on species diversity, and it was a good indicator of plant community diversity.

## INTRODUCTION

Ecological stoichiometry is an important tool for studying ecological processes and ecological functions(Zechmeister-Boltenstern S *et al*. 2016; Shang B *et al*. 2018), which is mainly used to explore the dynamic balance of various elements and their interactions (Sterner & Elser 2002; Moorthi S D *et al*. 2016). The C, N, and P concentrations are vital source elements in plants, and their changes in characteristics limit plant growth(Zeng Q *et al*. 2016; Sperfeld E *et al*. 2017). It can be used to judge the nutrient limitation of plants and to reveal the utilization strategies of plants for nutrients (Sterner & Elser 2002). At present, stoichiometry has been applied as a new ecological research tool to all levels, such as molecules, cells, individuals, communities and ecosystems (Elser J J *et al*. 2007; Elser J J *et al*. 2010; Yan Z B *et al*. 2016). With the changes of the global environment,terrestrial ecosystems have also undergone significant changes in the aspects of structure, process, function and vegetation distribution pattern (Freudenberger L *et al*. 2012). Among them, the role of community ecological stoichiometry has become a hot issue for scholars all around the world (Zechmeister B S et al. 2016; Bell D W *et al*. 2018; Yang Y *et al*. 2018.). The changes in plant ecological stoichiometry will contribute to nitrogen limitation, phosphorus limitation and both nitrogen and phosphorus limitations in the communities(Seastedt T R *et al*. 2001; Reich P B *et al*. 2006; Elser J J *et al*. 2007), which in turn affect community species diversity. It has been widely recognized that biological diversity plays an essential role in structuring communities and ecosystem processes (Tilman *et al*. 1996; Naeem & Li 1997; Yachi & Loreau 1999; Snyder *et al*. 2006; Haddad *et al*. 2011; Carlos Bustos-Segura *et al*. 2017). Community species diversity is a complex measure of community structure and function, and it is also the material basis for maintaining ecosystem stability and sustainable production(Vilela D S *et al*. 2016; Pelini S L *et al*. 2016; Zhang C H *et al*. 2017; Saitta A *et al*. 2018). Therefore, the use of ecological stoichiometry knowledge to study plant communities and their relationship to species diversity can help to reveal the ecological strategies of plants in specific environments, and is of great significance to the circulation and balance mechanism of community materials.

Understanding nutrient element utilization of plant has long been focus of plant ecology (Frost P C *et al*. 2012; Yan W *et al*. 2016). Since the 1980s, ecological stoichiometry has been applied to ecology for the first time(Reinhardt S B *et al*. 1986), it has become an important method for plant ecology research, and has been widely praised by ecologists all around the world(Allen A P. 2009; Castellanos A E *et al*. 2018; Yu H *et al*. 2017; Yan W *et al*. 2016). Most of the researches focuses on different ecosystem such as grassland ecosystem, forest ecosystem, wetland ecosystem and so on (Yang Y H *et al*. 2011; Cao Y *et al*. 2017; Grażyna P *et al*. 2018). And the rare researches that pay attention to desert ecosystems. Many researches contents is about the C, N, P concentrations and the ratios of plants at the organ and individual levels, which also get the results that C, N, P concentrations and their ratios of different organs were significantly different, the different individuals too (Zeng Q *et al*. 2017; Halvorson H M *et al*. 2017). However, in different ecosystems, the strategies for plant utilization of nutrients in different plant communities are not the same.

The C:N:P ratio is also considered to be one of the major drivers of biodiversity (Sardans J *et al*. 2011). For over a decade, evidence has accumulated suggesting a general trend towards the effect of community ecological stoichiometry on plant species diversity (Kerkhoff A *et al*. 2005; Liu Z *et al*. 2018). Importantly, this work suggests that community ecological stoichiometry can be important for plant species diversity (Huang J *et al*. 2018).The existing researches are mostly stoichiometry of plant communities with different functional groups and changes in species diversity in different successional stages.(Sardans J *et al*. 2011; Yan Z *et al*. 2015; Soufbaf M *et al*. 2018). However, the relationship between plant community diversity and ecological stoichiometry in desert systems has not been well-studied to our knowledge.

China is one of the most biodiverse countries on Earth(Chen & Bi 2007). Because northwestern China has been disturbed by climate change and by anthropogenic disturbances (Zhang Q *et al*. 2018), the plant communities of this region are extremely vulnerable and are expected to experience sharp reductions in population density in the future (Liu J *et al*. 2015). Plant communities in the upper reaches of the Tarim Desert are an especially fragile ecological zone (Gong Lu *et al*. 2016), but have been understudied. Hence, only a great understanding of the impact of ecology stoichiometry on plant species diversity is required to better know the reason of spatial distribution of plant speices, and in order to provide scientific basis for restructing and restoring the damaged ecological environment in the the upper reaches of the Tarim River. Here, we explore the following questions related to ecological stoichiometry and community diversity of this region: (1) what are the characteristic features of the community stoichiometry for this system? (2) to what extent does C, N, P contents of desert plants affect community diversity? The objectives of this study were to investigate whether desert plant community stoichiometry accounts for characteristics of trees, shrubs, and herbs, and to determine whether these relationships have any consequences for community diversity.

## MATERIALS AND METHODS

### Study area

Data were collected in the northern region of the Taklimakan Desert in Xinjiang, China (80°10’-84°36’E, 0°25’-41°10’N). The mean annual temperature of the area is approximately 10.4 °C, and the mean annual precipitation and evapotranspiration are roughly 50.4 mm, and 1800 mm, respectively (Gong L *et al*. 2016)The dominant plant species in this desert community are *Populus euphratica, Halostachys caspica, Tamarix chinensis, Phragmites communis, Lycium ruthenicum, and Glycyrrhiza uralensis*. The diversity of the natural vegetation of this area is low. The soil composition mainly is sand (Table. 1).

**Table 1.**
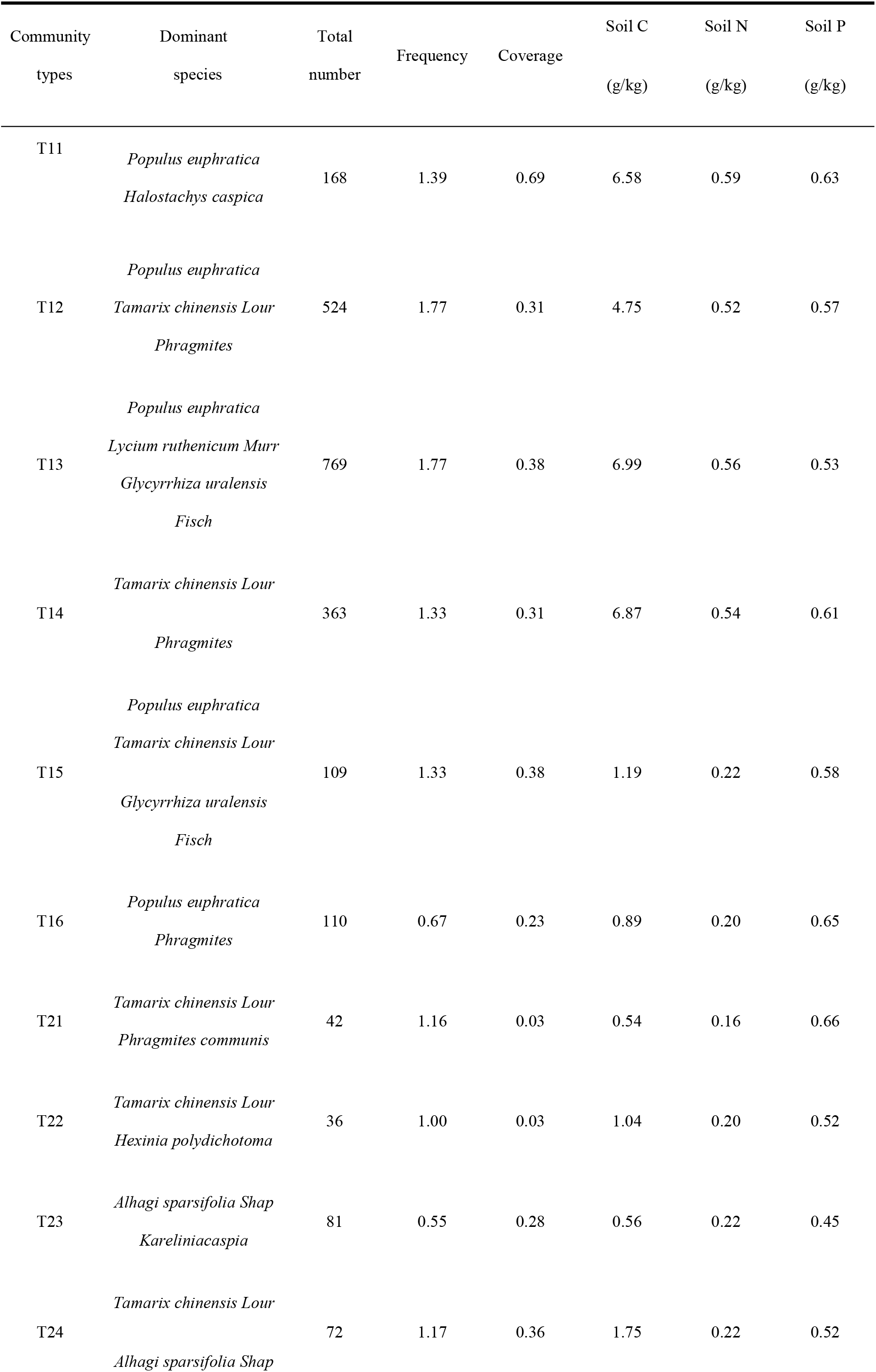
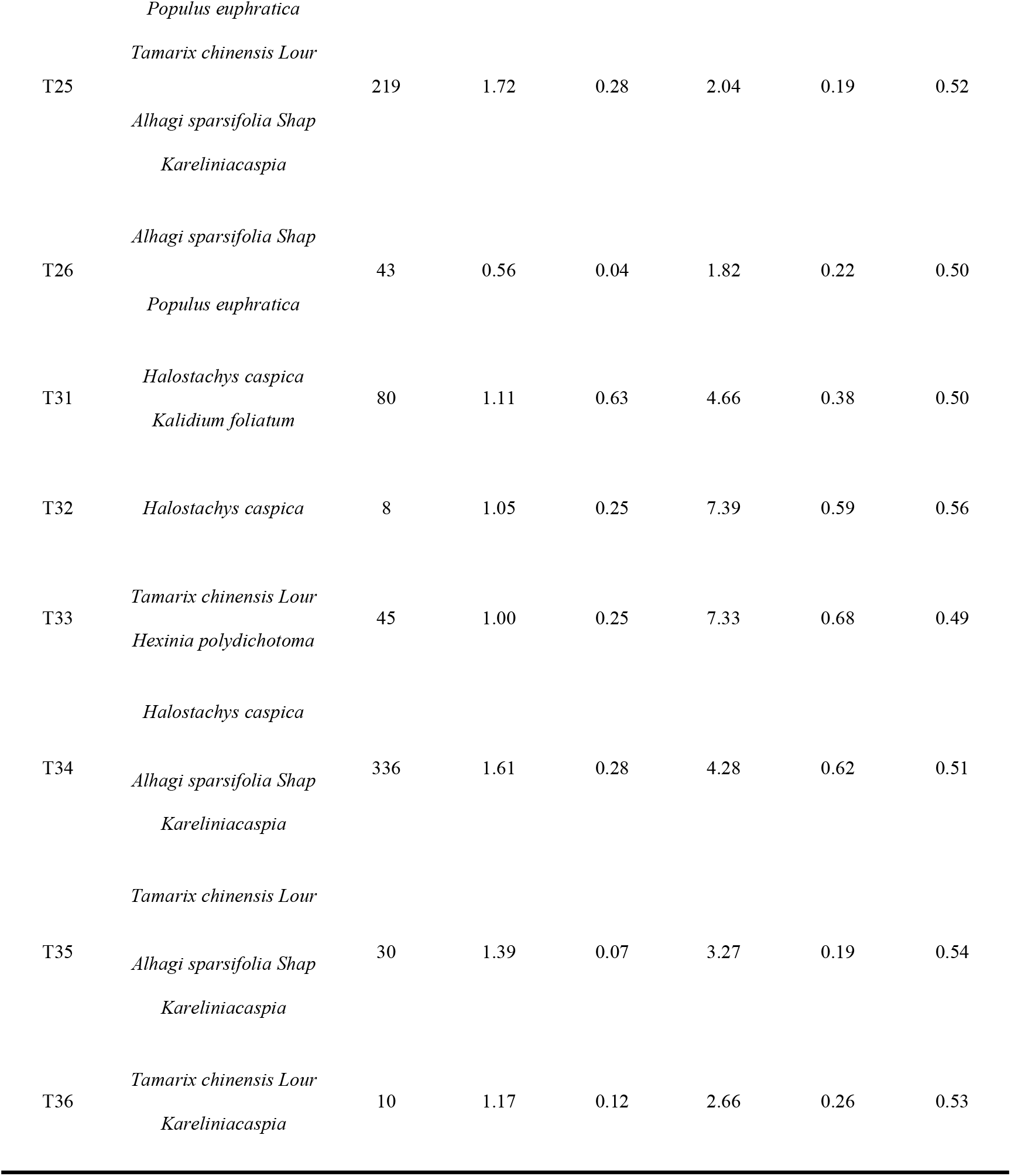
Vegetation composition and ecological characteristics of different community

### Field Methods

Data were collected at three separate sampling zones within the upper reaches of the Tarim River (Northwest China in August of 2016 (Figure 1). The three zones were selected so as to be in the same direction of the vertical river on the south side of the upper reaches of the Tarim River named T1,T2 and T3. According to the distance from the river, T1,T2 and T3 are in order. Within each of the three sampling zones, six individual plots were established to sample vegetation. The name rule about T1sample zone were T11,T12,T13,T14,T15 and T16. T2 sample zone and T3 sample zone naming rules were the same as T1 sample zone. Plots were spaced at intervals of at least 1 km within each sample zone, so as to avoid problems with pseudoreplication. This design resulted in a total of eighteen sampling plots. Plots were spaced at intervals of at least 1 km within each sample zone, so as to avoid problems with pseudoreplication. This design resulted in a total of eighteen sampling plots.

**Figure 1:**
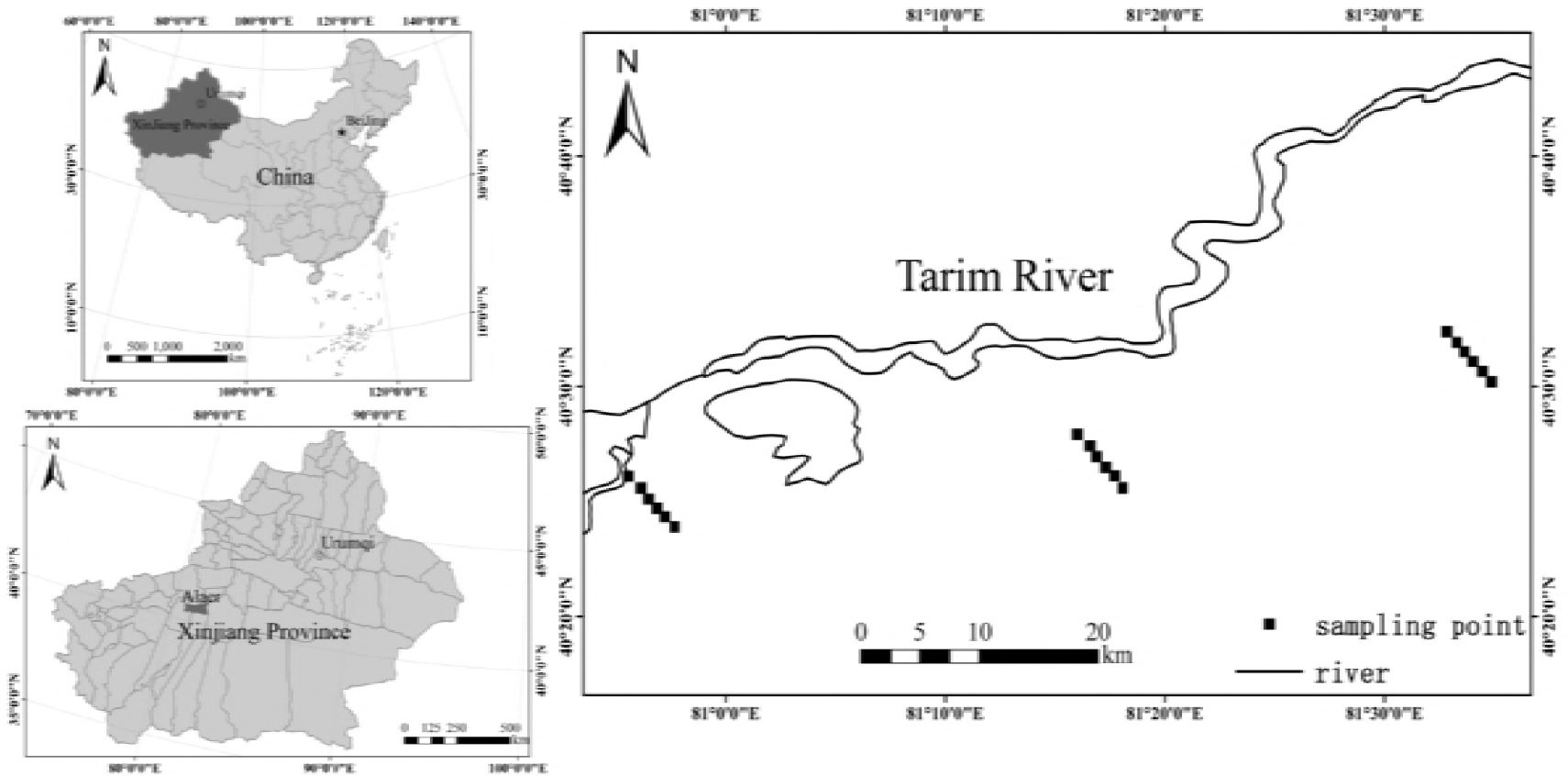
study area. *The picture in the upper left corner is the administrative division map of China, the dark Part is the Xinjiang Uygur Autonomous Region where the study area belongs, and the light part is the rest of the interior of China; The picture in the lower left corner is the administrative division map of Xinjiang Uygur Autonomous Region. The dark part is Alar where the administrative division belongs to the administrative division, and the light part is the other areas of Xinjiang; The picture on the left is the upper reaches of the Tarim River in the study area. The square is for each sampling point. The leftmost side is the TI sample with six community sampling points, the middle is the T2 sample with six community sampling points, and the rightmost is the T3 sample with six community sampling points.

Different plot sizes were used to sample trees, shrubs and herbs. Trees were sampled within 20 × 20 m^2^ plots, while shrubs and herbs were sampled within 10 × 10 m^2^ and 1 × 1 m^2^ plots, respectively. For trees, we recorded species names, tag numbers, plant height, crown width, diameter at breast diameter (DBH).For shrubs and herbs, we recorded species names, number, height, crown width, and total percent ground coverage. Photographs were taken of each individual for all plant species sampled. Healthy individuals were collected in the plot area. We collected approximately 50 g of leaf material from these individuals, and placed them in numbered envelopes with appropriate amount of desiccant. After removing litter and stones, we took the top soil (0–15 cm) of the plant crown using the quarter-division method.

### Experimental Analyses

Leaf samples were carefully cleaned and oven-dried at 60°C for two hours. Soil samples were air-dried after being sieved using a 2-mm mesh size, and visible roots and organic debris were removed by hand. All samples were then ground to a fine powder for element analysis (Soil Agrochemical Analysis (Bao S N, 2000). Total carbon content was determined by using the potassium dichromate external heating method (Bao S N, 2000), total nitrogen content was determined using the Kjeldahl method(Bao S N, 2000), and total phosphorus content was determined using the molybdenum-antimony colorimetric method (Bao S N, 2000). All analyses were repeated three times to take average.

### Community diversity indices

Next, we calculated several community diversity indices. These were as follows:

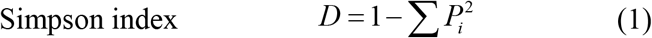

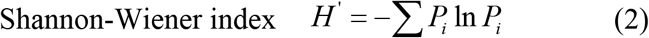

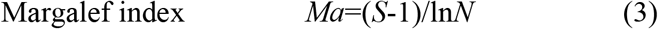

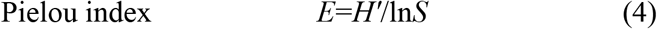

In the above equations, S is the total number of species in a plot, P_i_ is the ratio of the number of individuals in a species to the total number of individuals in a community. N is the total nuber of individuals in a plot.

### Community ecological stoichiometry

Leaf C,N, and P were used to calculate the importance values of each species in trees, shrubs and herbs. According to these importance values, the stoichiometric eigenvalues for all the species in the community were then weighted. In other words, the ecological stoichiometry of the community was the sum of the ecological stoichiometry of each weighted trees, shrubs or herbs (Yan E R, *et al*. 2008)

### Statistical Analyses

We used one-way analysis of variance (ANOVA) with a Duncan post hoc multiple comparisons test (Hsu J C. 1996.) to compare differences in C:N:P ratios among our communities.All statistical tests were assessed at a level of *α* = 0.05. Linear regressions were then performed to describe the relationships between C, N, P, C:N, C:P and N:P. This analyses were conducted in SPSS software (SPSS for windows. Version 19.0, Chicago, IL, U,S,A.)

Redundancy analysis (RDA) was then used to explore the correlation between each community index and community ecological stoichiometric concentrations and ratios. Structal equation model ( SEM) was constructed to examine the relationship between plant community stoichiometry and plant community diversity. RDA and SEM were conducted in CANOCO 4.5 software (Wright M G. 2014) and AMOS 24.0 software (SPSS for windows. Version 24.0, Chicago, IL, U,S,A).

## RESULTS

### The characteristic of plant community stoichiometry in different riparian zones

We found C concentration was greatest in T13 community in T1 zone and lowest in T31 community in T3 zone. Nitrogen concentration was greatest in T23 and was lowest in T14. Phosphorous concentration was greatest in T34 and was lowest in T22. All of the stoichiometric ratios (i.e., C:N, C:P and N:P) were greatest in zone 1 and were lowest in zone 3 (Table 2). Significant difference in the stoichiometric concentrations and stoichiometric ratios were observed among same zones. Significant differences in stoichiometry and stoichimetric ratios were also observed among the different zones. In addition, N and P concentrations were not significantly different among T15, T25 and T35 types (Table 2).

**Table 2.** The characteristic of C,N,P concentrations and ratios in different communities. *The lowercase letters are the differences in concentrations and ratios at different distances in the same vertical riparian, and the capital letters are the differences concentrations and ratios in the same distance at different vertical riparian.

| Community | C | N | P | C:N | C:P | N:P |
| --- | --- | --- | --- | --- | --- | --- |
| types | (mg/g) | (mg/g) | (mg/g) |  |  |  |
| T11 | 428.33 ± 1.04Bb | 16.74 ± 0.06Ad | 1.32 ± 0.07Aa | 26.05 ± 1.94Bbc | 326.04 ± 19.39Cc | 12.53 ± 0.34Bd |
| T12 | 447.70 ± 11.76Aa | 18.58 ± 0.96Aab | 1.10 ± 0.04Ab | 24.12 ± 0.62Bbc | 408.51 ± 9.94Cb | 16.94 ± 0.58Bc |
| T13 | 453.39±3.43Aa | 19.59±0.92Aa | 1.09±0.02Ab | 23.18±1.28Bc | 414.34±5.02ABb | 17.92±1.16Ac |
| T14 | 448.11±7.47Aa | 10.42±0.85Bd | 1.30±0.06Aa | 43.18±2.86Aa | 344.59±9.98Bbc | 7.99±0.33Be |
| T15 | 434.91±2.33Ab | 17.47±1.01Abc | 0.67±0.09Bc | 24.95±1.30Abc | 653.89±89.28Aa | 26.14±2.35ABa |
| T16 | 449.92±5.48Aa | 16.85±1.23Abc | 0.76±0.05Bc | 26.78±1.71Bb | 596.76±39.12Aa | 22.39±2.59Ab |
| T21 | 451.39±1.00Aa | 11.40±1.17Bb | 0.95±0.01Bc | 39.87±4.01Aa | 474.62±4.67Bcd | 11.99±1.30Bb |
| T22 | 420.51±1.05Bb | 14.64±1.04Bb | 0.59±0.01Ce | 28.82±1.20ABbc | 714.55±17.64Aa | 24.89±2.26Aa |
| T23 | 441.61±10.29ABa | 20.67±2.26Aa | 0.92±0.10Bc | 21.51±2.02Bd | 486.68±59.06Ac | 22.93±4.94Aa |
| T24 | 445.35±9.88Aa | 13.13±1.64Ab | 1.07±0.06Bb | 34.24±4.07Bab | 418.7±32.72Ad | 12.34±1.65Ab |
| T25 | 435.52±13.37Aab | 18.76±2.57Aa | 0.69±0.02ABd | 23.56±3.81Acd | 629.96±34.95ABb | 27.07±3.11Aa |
| T26 | 400.73±10.03Bc | 14.51±1.92ABb | 1.36±0.03Aa | 27.97±4.21Bbcd | 295.74±13.95Be | 10.70±1.33Bb |
| T31 | 391.05±1.46Cc | 11.35±0.84Bc | 0.69±0.01Cc | 34.58±2.66Aa | 562.76±4.96Aa | 16.07±0.73Abc |
| T32 | 405.37±5.95Bbc | 11.62±2.00Cbc | 0.71±0.02Bc | 35.67±6.84Aa | 567.44±16.64Ba | 16.05±0.72Bbc |
| T33 | 432.19±11.33Ba | 14.07±1.59Bbc | 1.26±0.11Ab | 35.22±4.88Aab | 345.24±24.51Bc | 11.31±2.23Bc |
| T34 | 433.15±22.08Aa | 14.58±0.02Ab | 1.46±0.10Aa | 29.72±1.54Bab | 297.79±6.26Cc | 10.05±0.72Bc |
| T35 | 419.32±11.91Aab | 17.93±2.81Aa | 0.83±0.08Ac | 23.80±3.93Ab | 507.29±52.38Bb | 21.50±2.34Ba |
| T36 | 443.39±18.88Aa | 12.84±0.45Bbc | 1.37±0.02Aab | 34.53±0.52Aa | 323.74±19.57Bc | 9.37±0.49Bc |

An extremely significant negative correlation but high slope was found between plant community C:N ratio and N concentration, and between C:N ratio and N:P ratio (Fig. 2A, 2D). Nitrogen concentration was not correlated with P concentration, although a strongly positively correlation emerged between N:P and N concentration (Fig. 2B). A significant negative correlation was found between plant community C:P ratio and P concentration (Fig. 2C). A significant positive was found between plant community C:P ratio and N:P ratio (Fig. 2E). There were no significant correlations between other concentrations and ratios.

**Figure 2:**
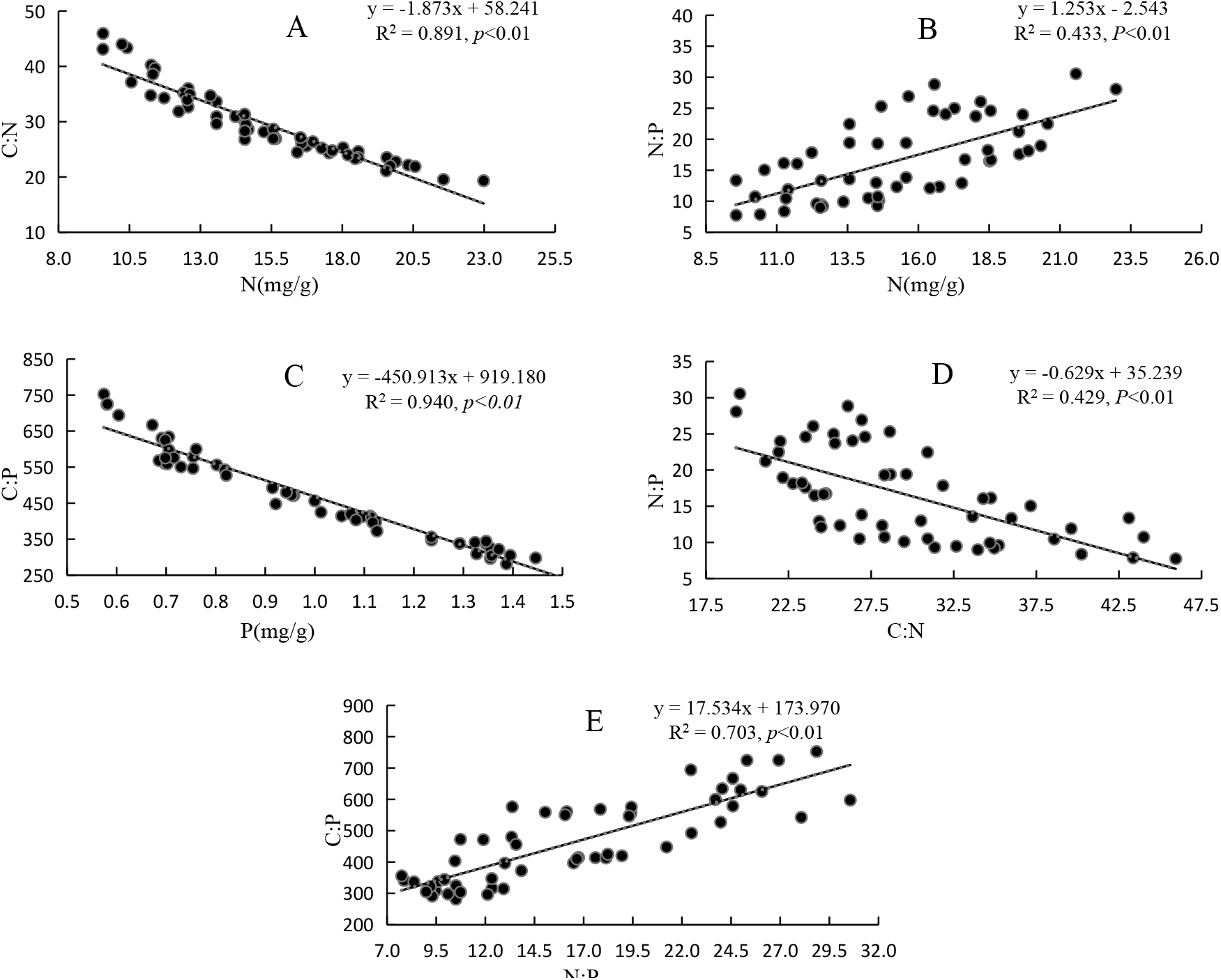
The correlation between communities C,N,P concentrations and ratio. *Fig 2 is a correlation diagram between stoichiometry concentrations and ratios. Where R^2^ represents the degree of fit, the closer R^2^ is to 1, the better the fit. *P* represents the correlation, *P* < 0.05 represents a significant correlation, *P* < 0.01 represents a extremely significant correlation; The equation is the Linear regression equation.

### The characteristic of plant community diversity in different ripirian zones

The Simpson index was highest in the T13 community (0.623, Fig. 3), and was lowest value in T31 (0.049, Fig. 3). The Shannon-Wiener index of the T11 and T16 communities was large, while the Shannon-Wiener index of the T31 community was smaller (Fig. 3). The Margalef index ranged from 0.227 in T23 to 0.64 in T12 (Fig. 3). The T23 and T11 communities had higher Pielou’s index values than the other communities (Fig. 3). Overall, the same general trends observed in the T1 zone plant community diversity incies generally greater than T2 and T3 plant community indices.

**Figure 3:**
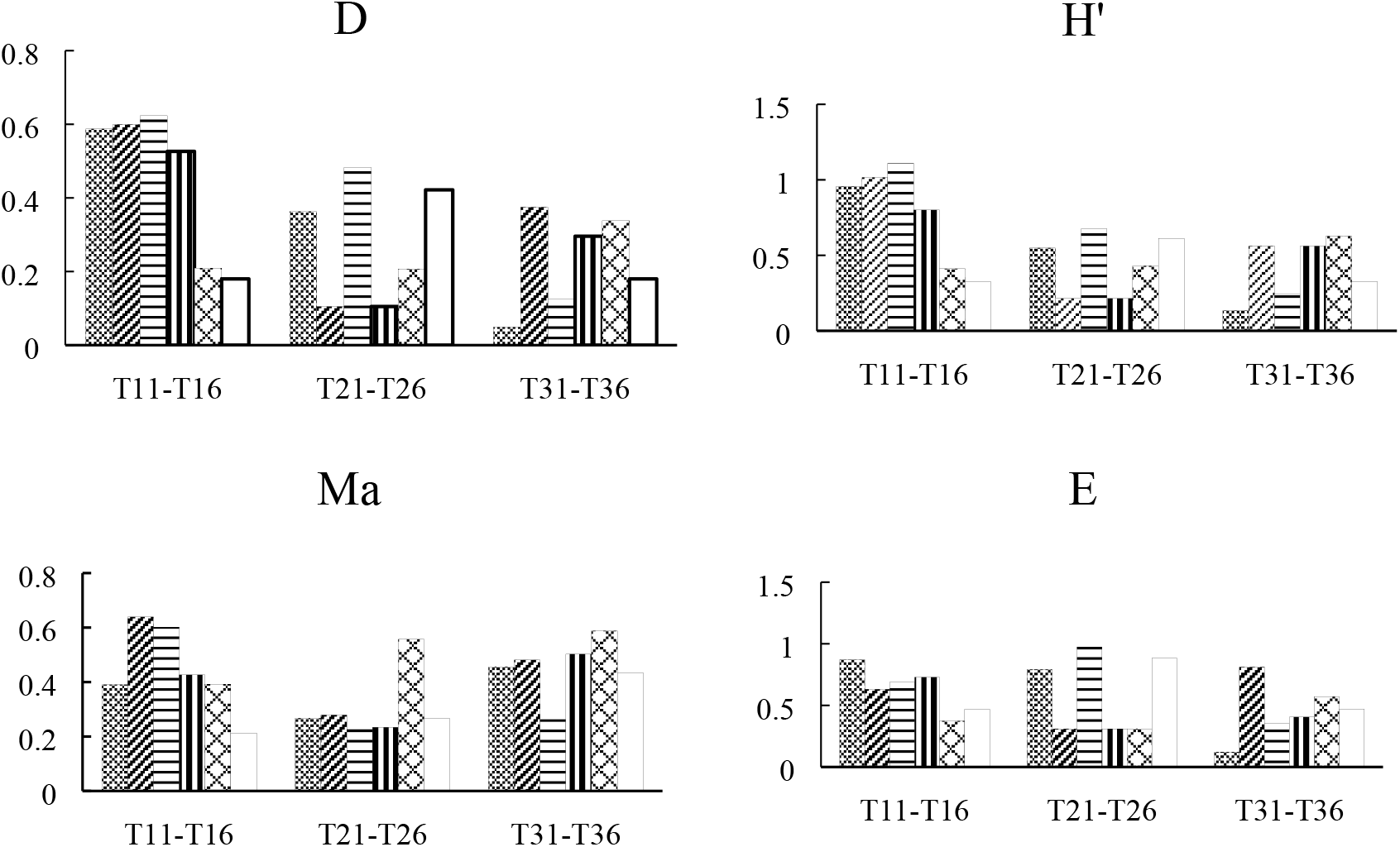
α diversity index in different plant community. *The bar graphs show that the values of *α* diversity indices, which are three ripirian zones consists of 18 communities.

### The correlation between plant community stoichiometry and plant community diversity

We used RDA to examine community diversity in more detail. The first axis explained 64% of the variation, while the second axis only explained 2%. As such, only axis 1 was a good reflection of the relationship between plant community stoichiometry and plant community diversity. In these biplots longer lines indicate stronger explanations, and smaller anglses indicate stronger correlations. The RDA indicated that the C:N ratio and N concentration played an important role in explaining plant community diversity among all concentrations and ratios. The Simpson’s index was strongly positively correlated with P concentration, and was weakly positively correlation with C and N concentrations (Fig. 4). In contrast, the C:N and N:P ratios had a strong negative correlation with Simpson’s index, and the C:P ratio showed a weak negative correlation (Fig. 4). A clear positive correlation emerged between the Shannon-Wiener index and C, N and P concentrations (Fig. 4). Likewise, a negative correlation among these ratios was found, and the remaining indices showed these simliar patterns as well (Fig. 4).

**Figure 4:**
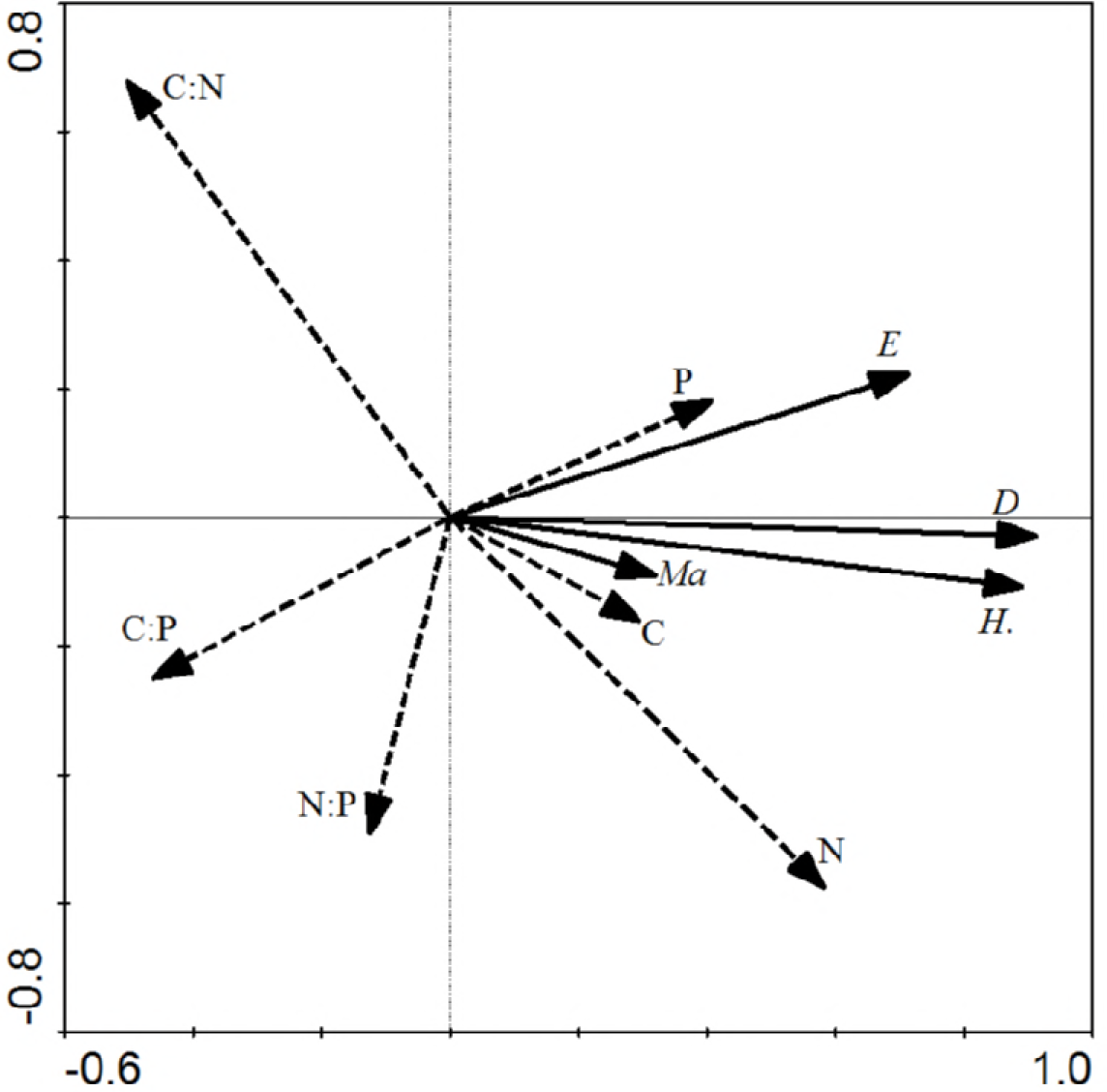
Stoichiometry and *α* diversity indices biplot diagram from the RDA. *The solid line indicate the community species diversity indices, and the dotted line indicate the community stoichiometry. In these biplots longer lines indicate stronger explanations, and smaller lines indicate weaker explanations. The angle indicates the correlation, the smaller the angle indicate the stronger correlation. When the angle is between 0° and 90°, there is a positive correlation between the two variables; when the angle is between 90° and 180°, there is a negative correlation between the two variables; when the angle is 90°, it means There is no correlation between the two variable.

Based on the above studies, it can be seen that the effects of concentrations and ratios on plant community diversity index were different, but it is not possible to conclude that the concentration or ratio had the greatest impact on plant community diversity index. Therefore, we performed a Monte-Carlo test on three concentrations and three ratios, and obtained the order of importance of the stoichiometric variables. The results are shown in Table 3. The important sequencing indicated that the most important element was N concentration (Table 3). We found that the N:P ratio was not as important for plant community diversity indices (Table 3). In order to further explore how C and N concentrations and C:N and C:P ratios potentially affect plant community diversity indices, we conducted a correlation analysis between single concentrations or ratios and plant community diversity indices. N, P concentrations and C:N, C:P ratios had great effect to four diversity indices. So to further explore the effects of these four important factors on the single plant community diversity index, We conducted a test of the impact of key concentration or ratio on the diversity index.The t-value biplot graph, if it falls into the circle with solid line, it means there is a significant positive correlation, and if it falls into the circle with dotted line it is a negative correlation. If the arrow line of the variable does not fall into the circle, indicating that the response variable has no correlation with this explanatory variable (Wright M G. 2014). Simpson’s index, Pielou’s index and the Shannon-Wiener index all fell within the real circle, and Margalef’s index partly fell into the circle with solid line (Fig. 5A). This shows that N concentration was strongly positively correlation with Pielou’s index, Simpson’s index and the Shannon-Wiener index, and was weakly positively with Margalef’s index. In other words, with an increase in N concentration, all of the indices become large. The C:N ratio revealed similar patterns as N concentration (Fig. 5C). Simpson’s index and Shannon-Wiener index fell completely into the the circle with solid line, and Pielou’s index and Margalef’s index partly fell intothe circle with solid line (Fig. 5B). This explained that N concentration was strongly positively correlated with the Simpson’s index, Pielou’s index and Shannon-Wiener index, and was weakly positively correlated with Pielou’s index and Margalef’s index. The C:N ratio revealed similar patterns to those for C:P concentrations (Fig. 5D).

**Figure 5:**
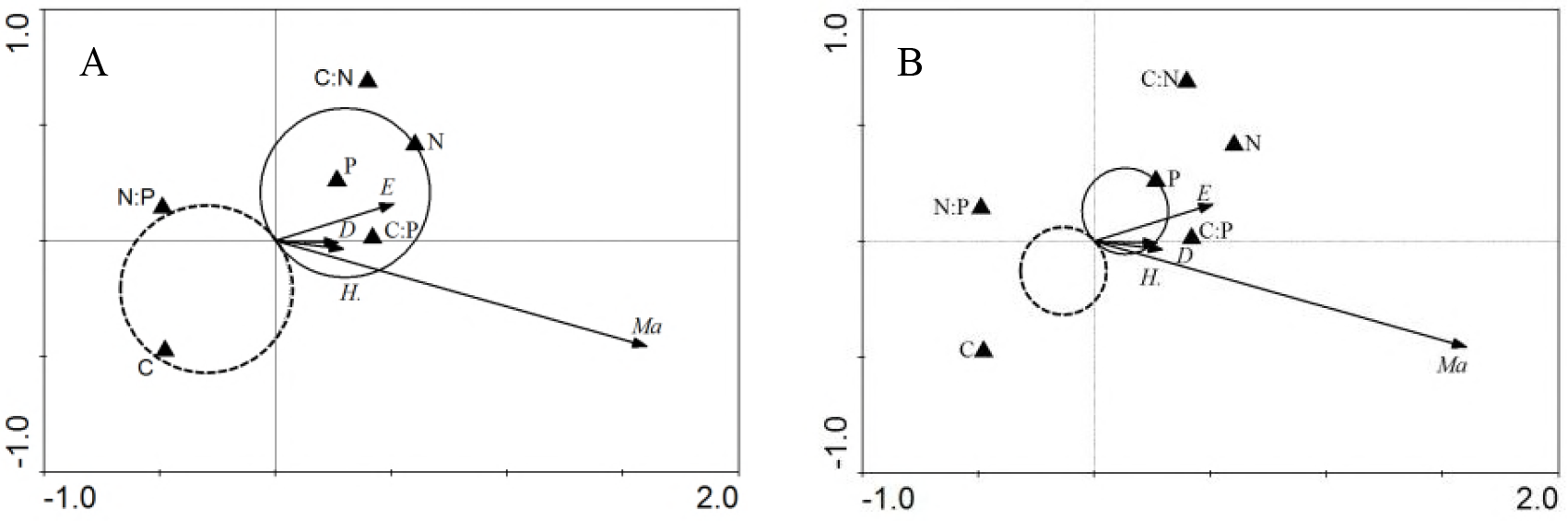
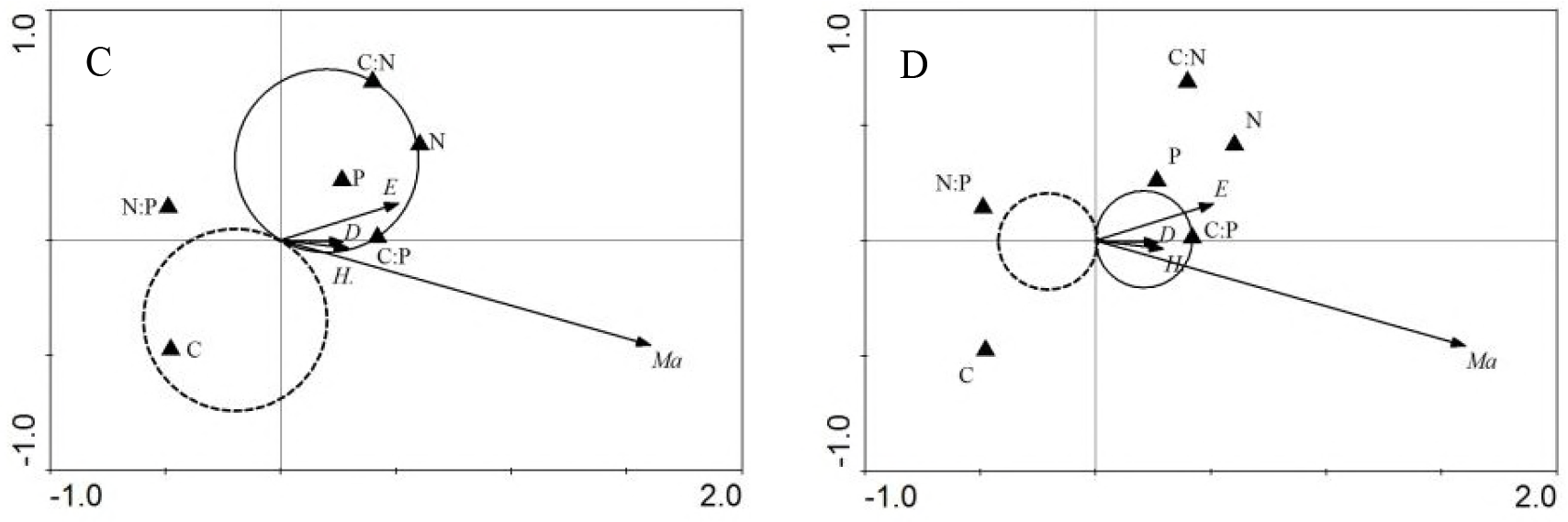
The t-value results for single factor influencing plant community diversity indices from RDA. *The solid circle indicates positive correlation, and the dotted circle indicates negative correlation. The α diversity indices fall into the solid circle, which indicate they have a positive correlation with the element or ratio. if they fall into the solid completely, that indicate they have a significantly positive correlation with the concentration or ratio; the α diversity indices fall into the dotted circle indicating a negative correlation with the concentration or ratio. The dotted circle indicates a significantly negative correlation if they fall into totally.

**Table 3.** Importance sequence and Duncan test of C,N,P concentrations and ratios from RDA

| Element | Importance sequence | Explanation | *F* | *P* |
| --- | --- | --- | --- | --- |
| N | 1 | 0.624 | 14.632 | 0.002 |
| C:N | 2 | 0.571 | 13.302 | 0.002 |
| C:P | 3 | 0.539 | 12.583 | 0.002 |
| P | 4 | 0.498 | 11.941 | 0.002 |
| C | 5 | 0.258 | 4.532 | 0.041 |
| N:P | 6 | 0.103 | 1.524 | 0.235 |

Structural equation models (SEM) were used to examine the relationship between plant community stoichiometry and plant community diversity. These results suggest that plant community stoichiometry was weakly positively correlated with plant community diversity (Fig. 6).

**Figure 6:**
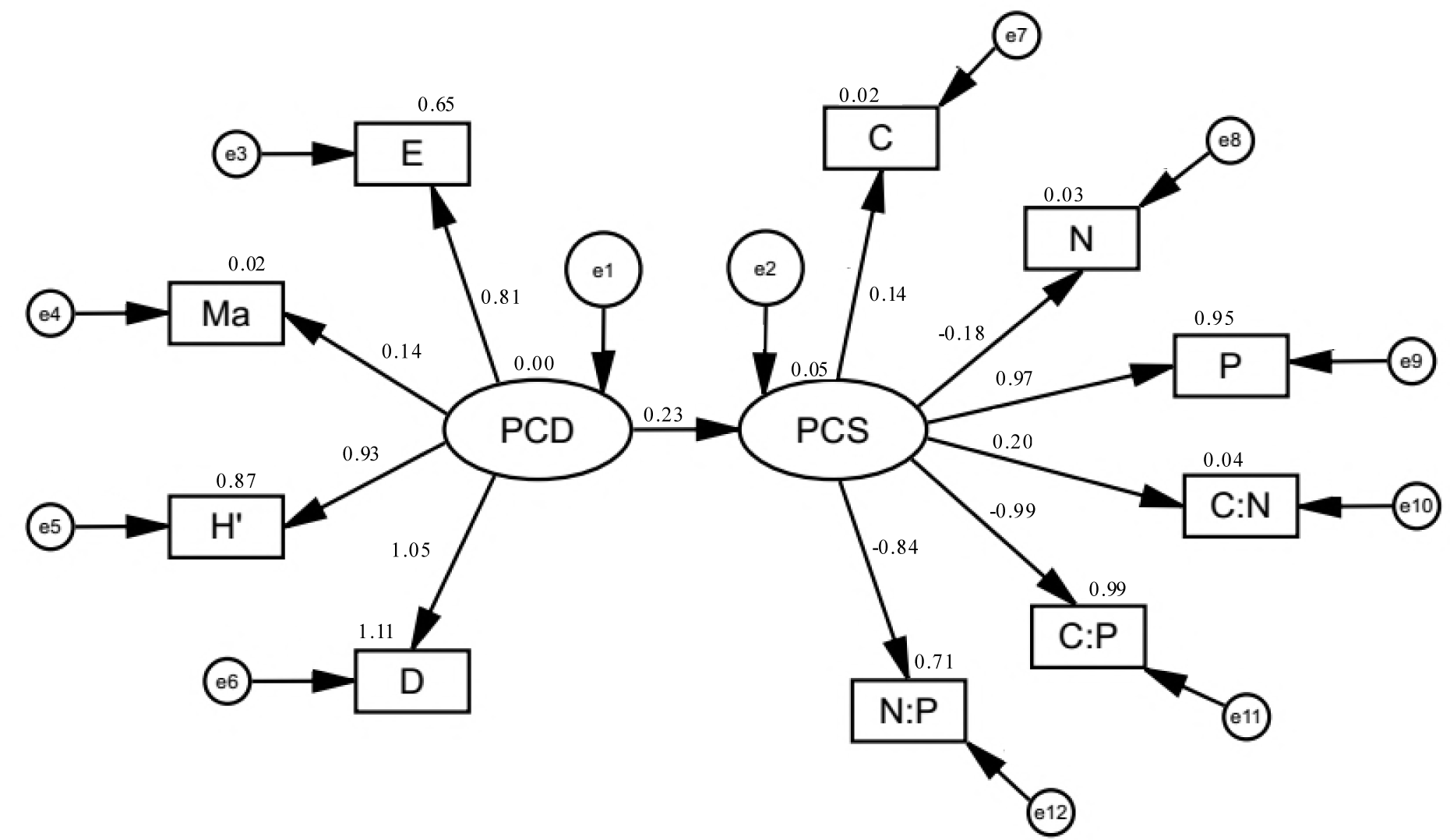
Structural equation models (SEM) of Plant community stoichiometry (PCS) and Plant community diversity (PCD) *The SEM of PCS and PCD, we used four α diversity indices to characterize PCS, at the same time 3 stoichiometry concentrations and 3 stoichiometry ratios are regard as PCD. Lowercase letter e is the error, and Number is the correlation index.

## DISCUSSION

### The effect of vertical riparian distance on community ecological stoichiometry

Our results demonstrate that C, N, and P concentrations in desert ecosystems is largely consistent with those from farmland ecosystems. These values may result from influences from oasis farmlands in the upper reaches of the Tarim River (Elser J J *et al*. 2007; Qu F *et al*. 2017). Indeed, our study area was adjacent to the Alar area. We observed large differences in C, N and P concentrations among different plant communities, and similar trends have also been reported for other areas (Behera S K *et al*. 2012). Carbon concentrations were highest at T13 in T1 zone. This community has the largest number of *Phragmites spp*, which is a unique xerophilous plant in the study area (Tuboi C *et al*. 2018). *Phragmites spp* are arid plants that like water, The T1 zone is closer to the water source than T2 zone and T3 zone, so a large amount of *Phragmites spp* grows in there. With the rapid growth of this species, the vascular bundles increase in the leaves and photosynthesis is enhanced, which may lead to the effective accumulation of organic matter in the leaves (Taysumov M A *et al*. 2017). The highest N concentrations were observed in the species of *Alhagi sparsifolia Shap*., which dominates the T23 community in T2 zone. *Alhagi sparsifolia Shap* is a naturally-growing and drought-tolerant plant, which absorbs groundwater and nutrients from the depths of the desert(Xiang Y L *et al*. 2018). If there is close to the water source, actually rich water, that is bad for *Alhagi sparsifolia Shap*, because it is a Xerophytes(Zhang B *et al*. 2018). On the contrary, there is far away from the water source, less water, can’t meet the plant own growth needs. So the right amount of water is most beneficial for the growth of the plant. *Alhagi sparsifolia* Shap is a leguminous plant that has strong N fixing ability and utilization efficiency (Lei L *et al*. 2014). The T11,T12,T13,T14,T15 and T16 communities in T1 zone, contained a high abundance of *Tamarix chinensis* Lour and its P concentrations are relatively high. The leaves of *Tamarix chinensis* Lour are scaly and contain higher P concentrations than most other species. This is due to the special healing body of stem and leaf that can enhance photosynthesis and produce a proteinaceous substance that provides a protective membrane to resist harsh environmental conditions (Rong Q *et al*. 2015). The P concentrations of the *Tamarix chinensis Lour* is the highest of any species. All in all, the traits of plant leaves appear to be a reflection of the characteristics of the plants themselves, which further reflects the specificity of the absorption and utilization efficiency of elements involved in the process.

In general, N and P are assumed to be the two most limiting elements to plant growth (Elser *et al*. 2000). We found a significant difference in N:P ratios among plant communities. Specifically, the N:P ratio were greater than 16 in some communities, indicating that these communities were strongly P-limited, and that other communities were more limited by N. In contrast, the plants in the upper reaches of the Tarim River are rarely restricted by N and P. A significant correlation between N concentration and C:N ratio in plant communities has been observed in other ecosystems (Yan Z B *et al*. 2015; Yu H *et al*. 2015). Even in desert ecosystems, C and N concentrations maintain a strong intrinsic connection, which is important for plants. On the other hand, P concentration and C:P ratio also showed a significant correlation, and a similar situation was observed in other areas as well (Plach J M *et al*. 2015). These results indicate that the C:P ratio of communities in arid regions is also limited primarily by P concentration.

### The effect of vertical riparian distance on community species diversity

The Simpson’s index, Shannon-Wiener index, Margalef’s index and Pielou’s index all serve as indicators of community diversity. Our result showed that the communities of T1 zones had the highest Simpson’s index values, indicating that they had strong anti-interference abilities. We believe that because of the river, it has been intermittently eroded by the Tarim River for a long time, making it more resistant to interference (Sures B *et al*. 1999; Bu C *et al*. 2017). However, other communities with poor anti-interference abilities have higher self-healing abilities after habitat destruction. This also indicates dominant species can play a leading role in maintaining community structure and ecological function (Zhang J *et al*. 2017).The Shannon-Wiener index of the T11 and T16 communities in T1 zone were large, indicating that the structure and composition of these communities are more complicated. In contrast, the T31 community in T3 zone had a low value, and was this relatively simple. The T11 and T16 communities in T1 zone are located closer to the Tarim River Basin and are not subject to drought stress. The water resources are relatively abundant and thus more plant species are capable of surviving there (Merritt D M *et al*. 2010). The T31 community in T3 zone is farther from the river, and only a handful of desert species that overcome drought stress have the ability to survive there. Pielou’s index can reflect the distribution of species in the community, where T23 and T11had greater evenness.

### The Relationship between plant community stoichiometry and plant community diversity

We found support for our hypothesis that plant community stoichiometry and plant community diversity were related. For example, each diversity index was significantly correlated with each stoichiometric concentration and ratio. We would this expect the stoichiometry of the community to be important at driving community diversity. Previous studies have found that C is the main element for plant, and compared with other nutrient concentrations, its concentration is high and stable (Heyburn J *et al*. 2017). However, C is not a key element that limits plant communities (Makino W *et al*. 2003; Heyburn J *et al*. 2007). Our results showed no significant correlation between plant community diversity and plant community C concentrations, further confirming this view. In this study, N and P were important to the plant community indices, and all showed a significant correlation with the N and P concentrations. This explains why changes in N and P concentration had an impact on plant community diversity (Zhao Y *et al*. 2010). Both the increase and decrease of N and P concentrations will affect the growth rate of plants, which will then cause changes in plant community diversity (Frenken T *et al*. 2017). The order of importance of community stoichiometry to community diversity shows that C:N and C:P ratios had significant impacts on community diversity. C:N and C:P ratios provide insight into plant’s ability to absorb nutrients and assimilate carbon (Bell D W *et al*. 2018). To a certain extent, it can also reflect the nutritional efficiency of the plant (Liu Z *et al*. 2010). Different communities have different carbon sequestration efficiencies, while carbon accumulation efficiency and storage capacities are associated with N and P supplies that limit plant growth. Ultimately, our results show that plant growth and community diversity are closely linked (Castellanos A E *et al*. 2018).

The SEM model also supported the notion that plant community stoichimometry is related to plant community diversity. The SEM was important at showing that these relationships are complex, and that no single plant community index or single concentration/ratio is apparent. The reason why the ecological stoichiometry characteristics can affect plant community diversity is due to Each plant community has different adaptation strategies for each element. (Fan H *et al*. 2015). On the one hand, it may be that the plants themselves have different preferences for the elements, and different plant communities are formed, resulting in different plant diversity. On the other hand, owing to the restriction of N and P elements in the local soil, the diversity characteristics of plant communities change, and the diversity of plant communities also change.

## ACKNOWLEDGEMENTS

The authors want to thank to the financial support of the National Natural Science Foundation of China (41461105). Additionally, we should also like to thank Dr. Emily Drummond at the University of British Columbia for her assistance with English language and grammatical editing of the manuscript.

